# Basolateral amygdala to nucleus accumbens projections differentially mediate flexibility of sign- and goal-tracking rats

**DOI:** 10.1101/2020.07.20.212738

**Authors:** Daniel E. Kochli, Sara E. Keefer, Utsav Gyawali, Donna J Calu

**Affiliations:** Department of Anatomy and Neurobiology, University of Maryland School of Medicine, 20 Penn Street, HSFII Room 203, Baltimore, MD 21201, USA; Program in Neuroscience, University of Maryland School of Medicine, Baltimore, MD 21201, USA

**Keywords:** Basolateral Amygdala, Nucleus Accumbens, Flexibility, Sign-tracking, Goal-tracking, Devaluation

## Abstract

Rats rely on communication between basolateral amygdala (BLA) and nucleus accumbens (NAc) to express lever directed approach in a Pavlovian lever autoshaping (PLA) task that distinguishes sign- and goal-tracking rats. While sign-tracking rats inflexibly respond to cues even after the associated outcome is devalued, goal-tracking rats flexibly suppress conditioned responding during outcome devaluation. Here, we sought to determine whether BLA-NAc communication in sign-trackers drives rigid appetitive approach that is insensitive to manipulations of outcome value. Using a contralateral chemogenetic inactivation design, we injected contralateral BLA and NAc core with inhibitory DREADD (hm4D-mcherry) or control (mcherry) constructs. To determine sign- and goal-tracking groups, we trained rats in five PLA sessions in which brief lever insertion predicts food pellet delivery. We sated rats on training pellets (devalued condition) or chow (valued condition) prior to systemic clozapine injections (0.1 mg/kg) to inactivate BLA and contralateral NAc during two outcome devaluation probe tests, in which we measured lever and foodcup approach. Contralateral BLA-NAc chemogenetic inactivation promoted flexible lever approach in sign-tracking rats, but disrupted flexible food-cup approach in goal-tracking rats. Consistent with a prior BLA-NAc disconnection lesion study, we find contralateral chemogenetic inactivation of BLA and NAc core reduces lever, but not foodcup approach in PLA. Together these findings suggest rigid appetitive associative encoding in BLA-NAc of sign-tracking rats hinders the expression of flexible behavior when outcome value changes.

## 1 Introduction

A body of evidence suggests that sign- and goal-tracking differences predict vulnerability to Substance Use Disorder (SUD) (Tomie et al., 2008; Flagel et al., 2009; Saunders & Robinson, 2010; Saunders et al., 2013; Kawa et al., 2016; Yager et al., 2015; Villaruel & Chaudhri, 2016). Reward predictive cues acquire appetitive motivational properties; a psychological process often referred to as incentive salience that is postulated to drive SUD vulnerability (Berridge, 1996; Robinson & Berridge, 1993; Berridge & Robinson, 2016). Sign-tracking (ST) and goal-tracking (GT) individual differences during a Pavlovian lever autoshaping task capture the degree to which reward predictive cues acquire incentive salience (Flagel et al., 2009; Pitchers et al., 2015; Flagel & Robinson, 2017) and predict heightened drug-cue induced relapse despite negative consequences (Saunders & Robinson, 2010; Saunders et al., 2013). Prior to drug experience, ST rats inflexibly respond to cues after reward devaluation (Morrison et al., 2015; Nasser et al., 2015; Patitucci et al., 2016; Smedley & Smith, 2018; Keefer et al., 2020). A prior lesion study indicates that communication between the basolateral amygdala (BLA) and nucleus accumbens (NAc) is necessary for the acquisition and expression of lever approach that classifies ST rats (Chang et al., 2012). Here we aim to determine the extent to which the incentive salience process supported by BLA-NAc core communication interferes with the expression of flexibility in ST rats during outcome devaluation.

BLA and NAc are critically involved in Pavlovian incentive learning processes including second order conditioning (SOC) and outcome devaluation. SOC is a learning process that relies upon the positive incentive value of the conditioned stimulus (CS), while outcome devaluation relies upon the current value of the unconditioned stimulus (US) (Holland & Rescorla, 1975). Pre-training lesions of either BLA or NAc impair both SOC and outcome devaluation, while post-training lesions of BLA disrupt only outcome devaluation, but not SOC (Hatfield et al., 1996; Setlow, Gallagher, et al., 2002; Johnson et al., 2009; Singh et al., 2010). Instead, the expression of SOC is mediated by NAc (McDannald et al., 2013). Pre-training, contralateral lesions disconnecting the BLA and NAc impair both SOC (Setlow, Holland, et al., 2002) and lever approach (the approach response characterizing ST rats), while leaving intact food cup-directed behavior (the approach response characterizing GT rats) (Chang et al., 2012). Taken together, the BLA and NAc support incentive learning relying on both conditioned stimulus (CS) value and current outcome (US) value. A growing number of studies demonstrate that GT, but not ST, rats flexibly reduce approach after outcome devaluation induced by satiety or illness (Morrison et al., 2015; Nasser et al., 2015; Patitucci et al., 2016; Smedley & Smith, 2018; Rode et al., 2020; Keefer et al., 2020). Both ST and GT rats similarly acquire and express SOC (Nasser et al., 2015; Saddoris et al., 2016), suggesting sign- and goal-trackers may utilize underlying BLA-NAc circuitry to differentially mediate incentive learning relying on CS or US value. Given tracking-related behavioral differences in incentive salience processing and flexibility, we hypothesize that the BLA to NAc communication drives rigid CS approach in ST rats and outcome value sensitive behavior in GT rats.

The primary prediction of our hypothesis is that contralateral chemogenetic inactivation of BLA and NAc core will make ST rats more flexible in outcome devaluation. Specifically, in intact ST rats we expect similar levels of responding for valued and devalued conditions, consistent with our prior reports (Nasser et al., 2015, Keefer et al., 2020). However, with BLA-NAc inactivation we predict reduced lever-directed approach for devalued relative to valued conditions. We expressed inhibitory DREADDs in contralateral BLA and NAc core and use systemic injections of low-dose clozapine to inactivate these structures during outcome-specific satiety devaluation. Because of the unidirectional and predominately unilateral projections of BLA to NAc (Swanson & Cowan, 1975; Ottersen, 1980; Russchen & Price, 1984; Heimer et al., 1991; Brog et al., 1993; Kelley et al., 1993), contralateral inactivation of the these structures disrupts communication from BLA to NAc core, while leaving an intact BLA and NAc core to support behavior that relies on either of these structures alone.

## 2 Materials and Methods

### 2.1 Subjects and Apparatus

We maintained male and female Long-Evans rats (Charles River Laboratories, Wilmington, MA; 250-275 g at time of arrival) (N = 98) on a reverse 12 h light/dark cycle (lights off at 9:00 AM). We conducted all behavioral training and testing during the dark phase of the cycle. All rats had ad libitum access to water and standard laboratory chow before being individually housed after surgical procedures. After recovery, we food restricted rats and maintained them at ~90% of their baseline body weight throughout the experiment. We performed all experiments in accordance to the “Guide for the Care and Use of Laboratory Animals” (8th edition, 2011, US National Research Council) and were approved by the University of Maryland, School of Medicine Institutional Animal Care and Use Committee (IACUC).

Prior to any training, we performed intracranial viral injection surgeries to deliver AAV8.hSyn.hM4Di.mCherry (hM4Di) or AAV8.hSyn.mCherry (mCherry) targeting the BLA and contralateral NAc core. We excluded some rats from subsequent analyses due to poor health or misplaced viral expression based on histological analysis (Figure 4), resulting in 72 rats being included in our analyses. The PCA characterization completed after surgery for viral injections resulted in the following number of rats in each group: ST n = 20 (mCherry n = 9 (n = 5 female, n = 4 male), hM4Di n = 11 (n = 7 female, n = 4 male), GT = 22 (mCherry n = 10 (n = 4 female, n = 6 male), hM4Di n = 12 (n = 7 female, n = 5 male), and INT n = 28 (mCherry n = 18 (n = 10 female, n = 8 male), hM4Di n = 10 (n = 3 female, n = 7 male).

We conducted behavioral experiments in individual standard experimental chambers (25 × 27 × 30 cm; Med Associates) located outside of the colony room. Each chamber was housed in an individual sound-attenuating cubicle with a ventilation fan. During PLA and devaluation probe tests, each chamber had one red house light (6 W) located at the top of the wall that was illuminated for the duration of each session. The opposite wall of the chamber had a recessed foodcup (with photo beam detectors) located 2 cm above the grid floor. The foodcup had an attached programmed pellet dispenser to deliver 45 mg food pellets (catalog#1811155; Test Diet Purified Rodent Tablet (5TUL); protein 20.6%, fat 12.7%, carbohydrate 66.7%). One retractable lever was positioned on either side of the foodcup, counterbalanced between subjects, 6 cm above the floor. Sessions began with the illumination of the red house light and lasted ~26 minutes.

### 2.2 Surgical Procedures

We rapidly anesthetized rats with 5% isoflurane and maintained them at 2-3% isoflurane (Vetone, Boisie, ID) throughout the procedure. We maintained body temperature with a heating pad during the procedure. Prior to the first incision, we administered a subcutaneous injection of the analgesic carprofen (5mg/kg) and subdermal injection of the local anesthetic lidocaine (10mg/ml at incision site). We secured rats in the stereotaxic apparatus (model 900, David Kopf Instruments, Tujunga, CA) and leveled the skull by equating lambda and bregma in the dorsal ventral plane. We lowered 10 μl Hamilton syringes (Hamilton, Reno, NV) into the brain targeting the BLA and contralateral NAc core (counterbalanced) using the following coordinates: BLA: (AP −3.0 mm, ML ± 5.0 mm, DV −8.6 mm 0° from midline) NAc core: (AP +1.8 mm, ML ± 2.5 mm, DV −7.0 mm −6° from midline) relative to bregma skull surface (Paxinos & Watson, 2007). We delivered AAV8.hSyn.hM4Di.mCherry (hM4Di) or AAV8.hSyn.mCherry (mCherry) targeting the BLA and contralateral NAc core (Addgene, Watertown, MA) via a micropump (UltraMicroPump III, World Precision Instruments, Sarasota, FL) at a volume of 600 nL per site at a rate of 250 nL/minute. We left syringes in place for 10 minutes after the infusion ended to allow diffusion of the viral constructs prior to suturing incisions. After surgery, we placed the rats into a recovery cage on a heating pad until ambulatory. We administered Carprofen (5 mg/kg; s.c.) 24 and 48 hours post-surgery and monitored weights daily to confirm recovery.

### 2.3 Pavlovian Lever Autoshaping Training and Testing

We trained rats over five daily Pavlovian lever autoshaping sessions (approximately 26 minutes duration per session), which consisted of 25 reinforced lever conditioned stimulus (CS+) presentations occurring on a VI 60 s schedule (50-70s). Trials consisted of the insertion of a retractable lever (left or right, counterbalanced) for 10 s, after which the lever was retracted and two 45 mg food pellets were delivered to the foodcup, non-contingent on rat behavior. The sessions took place in darkness with a red house light that was illuminated for the duration of the session.

After acquisition, we performed two days of satiety-induced outcome devaluation testing. Prior to test sessions, we gave rats free homecage access to 30g of rat chow (valued condition) or the same food pellets delivered during training (devalued condition) in a pre-habituated ceramic ramekin (similar to Parkes & Balleine, 2013). Immediately following satiation, we gave systemic injections of 0.1 mg/kg clozapine i.p. (Tocris, Bristol, UK) dissolved in bacteriostatic saline prior to transport to the behavioral chambers (Gomez et al. 2017). We waited 30 min after injection to allow binding of the ligand to the DREADD receptors. Then we gave a PLA probe test (approximately 10 minutes duration) consisting of 10 non-reinforced lever presentations occurring on a VI60 s schedule (50-70s). Immediately following testing, we gave rats a 30 min choice test in which they could consume up to 10g each of rat chow or pellets in the homecage. Between each PLA test we gave rats a single reinforced lever autoshaping training session to track stability in Pavlovian behavior. The next day, we gave rats a second round of satiety devaluation, PLA probe, and choice tests while sated under the opposite condition (pellet or chow; order counterbalanced).

### 2.4 Measurements

During PLA acquisition and probe tests, we collected three behavioral measurements during the 10 s CS (lever) period. All behavioral measurements were automatically collected and scored via MED-PC computer software (Med Associates, Georgia, VT). For foodcup and lever contacts, we recorded the total number of contacts and latency to first contact for all sessions. On trials in which no contact occurred, we recorded a latency value of 10s. We calculated the lever or foodcup probabilities by dividing the number of trials that a lever or foodcup contact was made by total number of trials in the session.

The criterion used for behavioral characterization of sign- and goal-tracking phenotype was based on a Pavlovian Conditioned Approach (PCA) analysis (Meyer et al., 2012) determined by averaging PCA scores during training sessions four and five. The PCA score quantifies the variation between lever directed (sign-tracking) and foodcup directed (goal-tracking) behaviors. Each rat’s PCA score is the average of three difference score measures (each ranging from −1.0 to +1.0): (1) preference score, (2) latency score, and (3) probability score. The preference score is the number of lever presses during the CS, minus the foodcup pokes during the CS, divided by the sum of these two measures. The latency score is the average latency to make a foodcup poke during the CS, minus the latency to lever press during the CS, divided by the duration of the CS (10 s). The probability score is the probability to lever press, minus the probability to foodcup poke observed throughout the session. Sign-tracking PCA scores range from +0.33 to +1.0, goal-tracking PCA scores range from −0.33 to −1.0, and intermediate group PCA scores range from −0.32 to +0.32.

### 2.5 Histology

After completion of behavioral testing, we deeply anesthetized rats with isoflurane and transcardially perfused them with 100 ml of 0.1 M PBS followed by 400 ml 4% paraformaldehyde in 0.1 M sodium phosphate, pH 7.4. We removed brains and post-fixed them in 4% paraformaldehyde for two hours before transfer to a 30% sucrose 4% paraformaldehyde solution in 0.1 M sodium phosphate for 48 hours at 4°C. We then rapidly froze them via dry ice and stored them at −20°C until sectioning. We collected 50 μm coronal sections through the entire extent of the nucleus accumbens and amygdala via a cryostat (Lecia Microsystems). We mounted sections on slides and verified viral expression in BLA and NAc core using anatomical boundaries defined by Paxinos and Watson (Paxinos & Watson, 2007) using a confocal microscope. The observer was blind to the condition and behavior of each animal.

### 2.6 Experimental Design and Statistical Analysis

Data was analyzed using SPSS statistical software (IBM v.25) with mixed-design repeated-measures ANOVAs. Analyses included the within-subjects factors of Response (foodcup, lever) and Value (valued, devalued) and the between-subjects factors of Virus (mCherry, hM4Di), Tracking (ST, INT, GT), and Sex (female, male) as indicated in results section. Unplanned post-hoc tests used a Bonferroni correction. Training analyses include all tracking groups (ST, INT, GT). Devaluation analyses include ST and GT rats to test a priori hypotheses based on previously reported flexibility differences in these two tracking groups (Keefer et al., 2020; Nasser et al., 2015). Due to the importance of using both males and females in research (McCarthy et al., 2017; Miller et al., 2017; Shansky, 2019), we explore the possibility of sex-differences by reporting sex effect sizes (Miller et al., 2017). Sex effect sizes are expressed as Cohen’s *d* (*d* = (M1 − M2) / SDpooled), where M1 is mean of group 1, M2 is mean of group 2, and SDpooled = √ (s12 + s22) / 2, which is the pooled standard deviation of the two groups (Cohen, 1988). This approach allows us to interpret potential sex effects that aren’t appropriately powered for typical statistical analysis. We follow general guidance for interpreting effect sizes where small effect *d* = 0.2, medium effect *d* = 0.5, and large effect *d* = 0.8 or larger (Cohen, 1988), and note instances that future studies should be powered to explore sex as a biological variable.

## 3 Results

### 3.1 Acquisition of Pavlovian Lever Autoshaping

We trained rats for five days in Pavlovian Lever Autoshaping to determine tracking groups prior to outcome devaluation testing. We used a Pavlovian Conditioned Approach Index (Fig. 1A, see methods for calculation) that takes into account the number of lever and foodcup contacts (Fig. 1B-C), latency to contact, and probability of contact for both lever and foodcup. We analyzed the lever autoshaping training data using six separate mixed-design, repeated measures ANOVAs with the between-subjects factor of Tracking (ST, INT, GT) with the within-subjects factors of Session (1-5). In Table 1 we report main effects and interactions of these analyses. Notably, the critical Session × Tracking group interactions were significant for all six measures of conditioned responding (*F*s>12.713, *p*s<0.001). We analyzed terminal levels of lever and foodcup contacts on Session 5, using between-subject factors of Virus (mcherry, hm4di) and Tracking (ST, INT, GT) and found no Virus main effects nor Virus x Tracking interactions (Fig. 1D) indicating that behavior did not differ between viral conditions prior to test for any of the six lever autoshaping measures (*F*s<3.3, *p*s>0.05). This was also the case when only ST and GT rats were included in the terminal contact analysis (all *F*s<2.48, *p*s>0.05).

**Figure 1.**
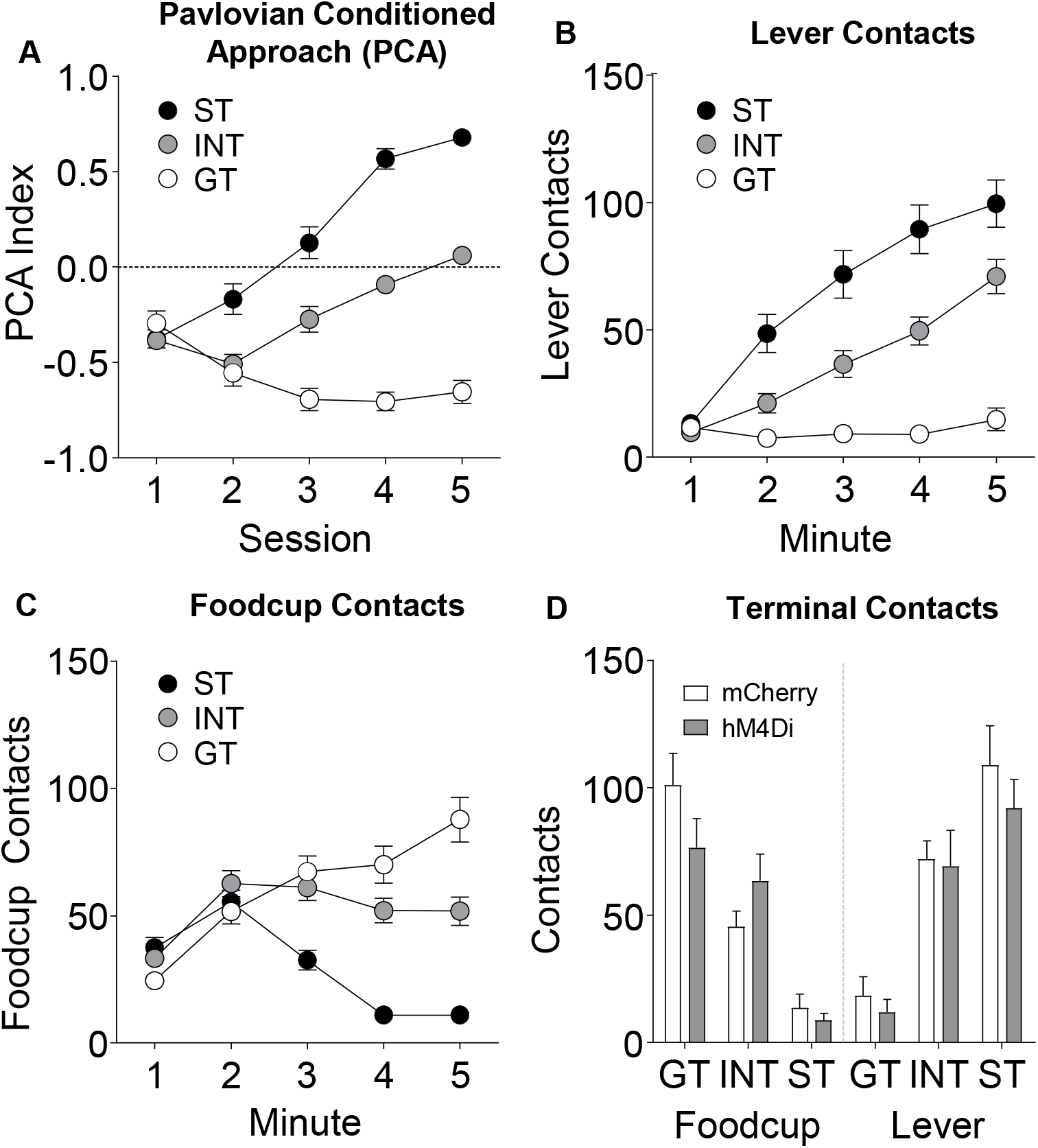
Pavlovian Lever Autoshaping acquisition data. Data represents (A) average PCA score, (B) lever contacts, (C) foodcup contacts during training; and (D) both terminal lever and foodcup contacts on fifth training session are represented as a function of viral condition.

**Table 1.** Repeated Measures ANOVA for Pavlovian lever autoshaping across all tracking groups

| Effect | Degrees of Freedom | Lever |  |  |  |  |  |
| --- | --- | --- | --- | --- | --- | --- | --- |
|  |  | Contact |  | Latency |  | Probability |  |
|  |  | *F* | *p* | *F* | *p* | *F* | *p* |
| Session | (4,268) | 59.805 | <.001 | 71.348 | <.001 | 72.357 | <.001 |
| Tracking | (2,67) | 43.386 | <.001 | 36.199 | <.001 | 49.196 | <.001 |
| Session * Tracking | (8,268) | 15.106 | <.001 | 12.713 | <.001 | 15.085 | <.001 |

| Effect | Degrees of Freedom | Foodcup |  |  |  |  |  |
| --- | --- | --- | --- | --- | --- | --- | --- |
|  |  | Contact |  | Latency |  | Probability |  |
|  |  | *F* | *p* | *F* | *p* | *F* | *p* |
| Session | (4,268) | 20.647 | <.001 | 36.887 | <.001 | 29.325 | <.001 |
| Tracking | (2,67) | 14.434 | <.001 | 27.219 | <.001 | 24.841 | <.001 |
| Session * Tracking | (8,268) | 25.267 | <.001 | 31.322 | <.001 | 28.135 | <.001 |

### 3.2 Effects of contralateral BLA-NAc core inactivation on Pavlovian approach during outcome devaluation

We hypothesized that ST rats rely on BLA-NAc core to drive rigid appetitive approach. To test this a priori hypothesis, we examined the extent to which BLA-NAc core contralateral chemogenetic inactivation altered the preferred response ST rats during satiety devaluation tests. For ST rats the preferred response is lever contacts (Fig. 2A), while for GT rats the preferred response is foodcup contacts (Fig. 2B). Notably, mCherry ST control rats showed no difference in lever contact between valued and devalued tests, confirming their insensitivity to devaluation, consistent with prior reports (Keefer et al., 2020; Nasser et al., 2015). ST rats expressing hm4di showed greater lever contact during valued compared to devalued tests (*t*(10)=2.582, *p*=0.027), indicating devaluation sensitivity in ST rats with contralateral chemogenetic inactivation of BLA-NAc core (Fig. 2A). In contrast, mCherry GT control rats showed greater foodcup contact during valued compared to devalued tests (*t*(9)=2.273 *p*=0.049), confirming their devaluation sensitivity that is consistent with prior reports (Keefer et al., 2020; Nasser et al., 2015). GT rats expressing hm4di constructs showed no difference in foodcup contact during valued compared to devalued tests, indicating contralateral chemogenetic inactivation of BLA-NAc core makes GT rats insensitive to devaluation (Fig. 2B). We also conducted a repeated measures ANOVA on these preferred response data using between-subjects factors of Virus (mCherry, hM4Di) and Tracking (GT, ST), and the within-subject factor of Value (valued, devalued). We observed main effects of Virus (*F*(1,38)=5.485, *p*=0.025) and Tracking (*F*(1,38)=42.461,*p*<0.001), as well as Value x Tracking (*F*(1,38)=4.552, *p*=0.039) and Virus x Tracking (*F*(1,38)=4.460, *p*=0.041) interactions (see Fig. 2 A-B), indicating both virus and value manipulations differ by tracking group. For parallel analyses of non-preferred responding (lever contact for GT and foodcup contact for ST rats), we observed a main effect of Tracking such that GT performed more non-preferred approach behavior, *F*(1,38)=7.773, *p*=0.008), but no other main effects or interactions, *p*s>0.05 (see Fig. 2 C-D).

**Figure 2.**
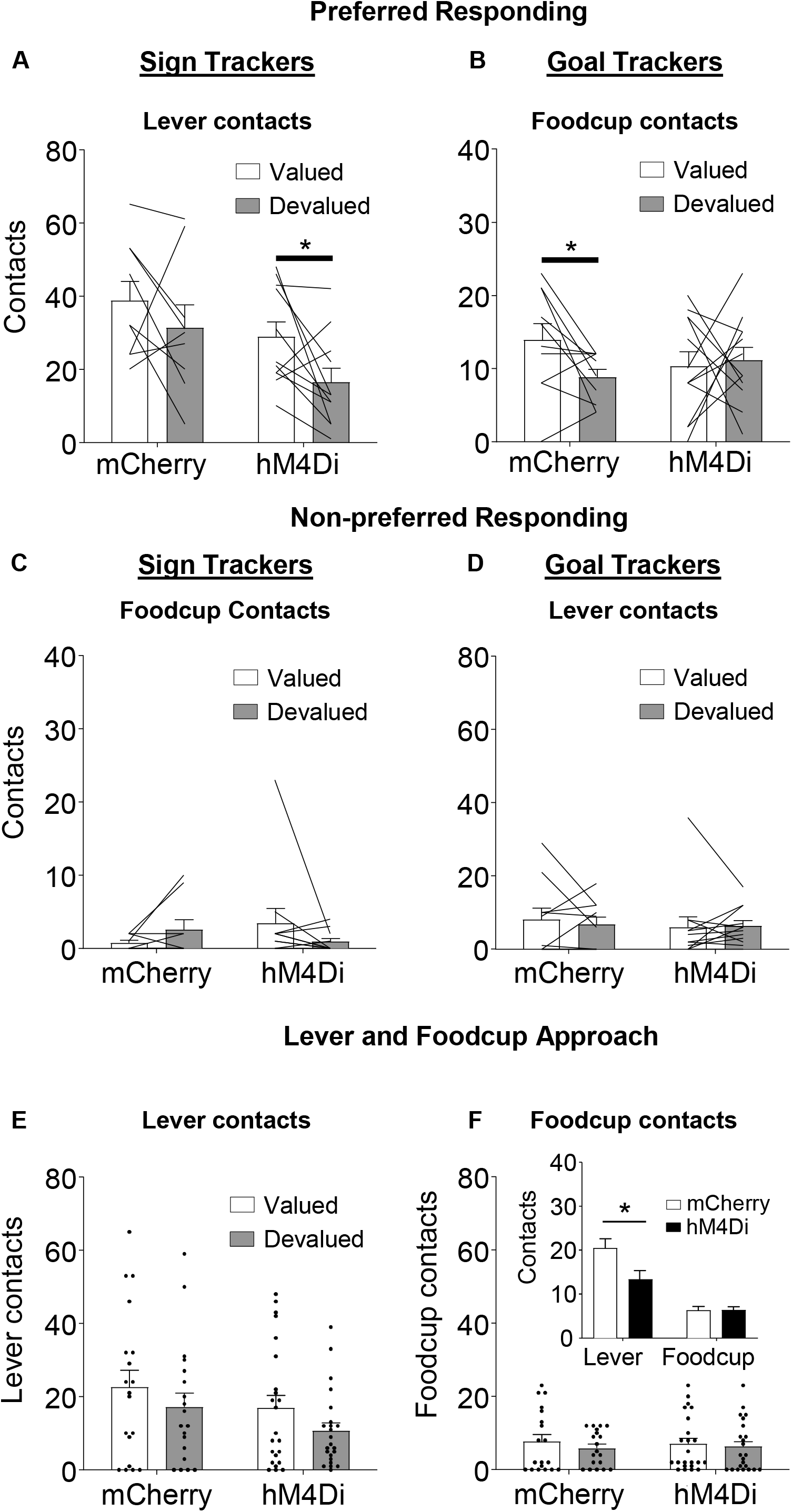
Outcome devaluation in sign- and goal-tracking rats. Data represents individual subjects (line) and group averaged (bars) for (A-B) preferred responding (ST: lever contact, GT: foodcup contact) and (C-D) non-preferred responding (ST: foodcup contact, GT: lever contact), + SEM. A priori planned comparisons reveal that (A) hM4Di, but not mCherry, ST show devaluation effect (difference between valued and devalued) for lever directed behavior, *t*(10)=2.582, *p*<0.05. (B) mCherry, but not hM4Di, GT show devaluation effect for foodcup directed behavior, *t*(9)=2.273 *p*<0.05. No differences were found for non-preferred responding. Data for (E-F) represents individual subjects (dot) and group averages (bars) for (E) lever and (F) foodcup contacts during outcome devaluation; (F inset) BLA-NAc core inactivation disrupts lever but not foodcup approach.

A prior lesion study demonstrated that BLA-NAc communication drives lever directed, but not foodcup directed behavior in lever autoshaping (Chang et al., 2012). To evaluate whether we replicate this BLA-NAc lesion finding using our contralateral inactivation approach, we analyzed the data by including Response (lever, foodcup) as a factor. Consistent with the prior study, we observed a Response x Virus interaction (*F*(1,34)=4.484, *p*=0.042), shown in Fig. 2E-F, in which lever approach is affected more by contralateral BLA-NAc core inactivation than foodcup approach across both value conditions (Fig. 2E-F and Fig.2F inset). Because we included both males and females in this study, we next examined whether Sex interacted with any other factors during our devaluation tests. In addition to main effects for all factors (Value, Response, Virus, Sex, and Tracking, all *F*>4.983, *p*<0.05), we also observed a Response x Sex interaction, *F*(1,34)=4.688, *p*=0.037), which we explore by separately analyzing each response.

We analyzed lever-directed behavior with between-subjects factors of Tracking (ST, GT), Virus (mCherry, hM4Di) and Sex (female, male), and within-subjects factor of Value (valued, devalued). Again, we observed a main effect of Sex (*F*(1,34)=5.549, *p*=0.024), driven primarily by more lever approach in females compared to males across virus groups and value conditions (Fig. 3A). We also observed main effects of Value (*F*(1,34)=8.527,*p*=0.006) and Virus (*F*(1,34)=6.114,*p*=0.019). We next analyzed foodcup-directed behavior using the same factors We observed a Value x Tracking x Sex interaction (Fig. 3B; *F*(1,34)=5.02, *p*=0.032).

**Figure 3.**
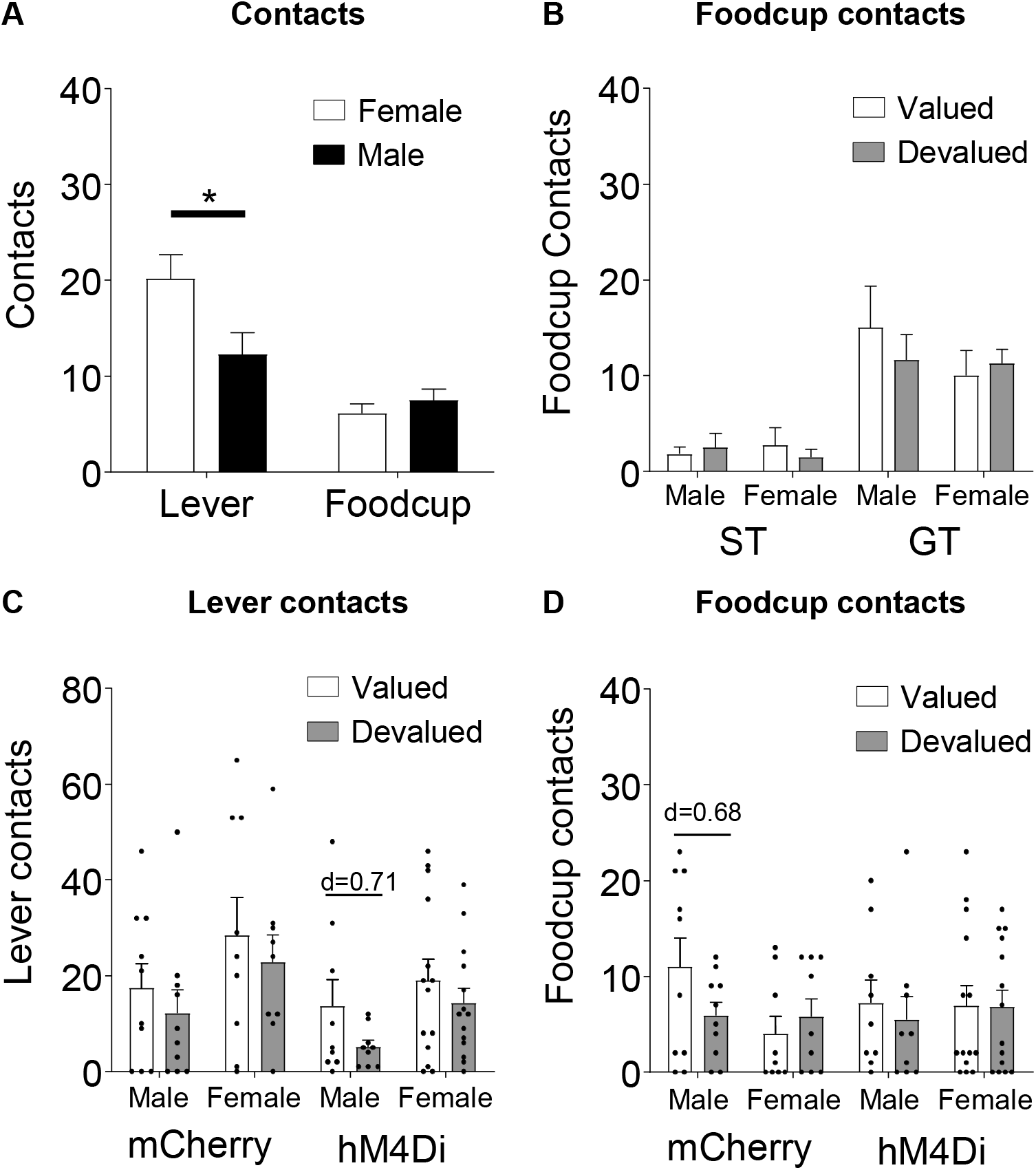
Sex effects during outcome devaluation; split by response type (A-B) and virus group (C-D). Data represents individual subjects (dot) and/or group averages (bars) + SEM. (A) Females preform more lever-directed responses than males during outcome devaluation tests overall. (B) Tracking x value x Sex interaction of foodcup responding. (C) Male hM4Di rats show moderate devaluation effect sizes for lever approach, Cohen’s *d*=0.71, whereas (D) intact mCherry males show moderate devaluation effect sizes for foodcup approach, Cohen’s *d*=0.69.

Finally, we show lever and food cup contact data for male and female rats within each viral group in Fig. 3C-D. We provide effect size calculations for transparent reporting of data from both sexes in our study of the effects of contralateral BLA-NAc core inactivation on lever and food cup approach in outcome devaluation. For lever-directed behavior, we observed a medium devaluation effect size only in hM4Di males (Cohen’s *d* = 0.71 valued vs. devalued), while small devaluation effect sizes were observed for male mCherry rats and females in both viral groups (Fig. 3C; Cohen’s *d* all < 0.33 valued vs. devalued). For foodcup behavior, we observed medium devaluation effect sizes only in males with BLA-NAc core intact (mCherry males Cohen’s *d* = 0.68 valued vs. devalued), while small devaluation effect sizes were observed for male hM4Di rats and females in both viral groups (Fig. 3D; Cohen’s *d*s all < 0.24 valued vs. devalued). These data suggest future studies designed to probe sex-specificity of BLA-NAc core manipulations may be warranted.

### 3.3 Satiety and Devaluation Choice Test

We recorded pellet and chow consumption during satiety (pre-test) and choice test (post-test). Prior to devaluation test sessions, we found no difference in the amount of food consumed between tracking or viral groups during the satiation hour (*F*<1, *p*>0.4). To confirm the devaluation of the sated food, we gave rats a post-satiety choice test following the devaluation test. Rats preferred to consume food they were not sated on, as indicated by a main effect of Choice, *F*(1,40)=46.125, *p*<0.0001. There were no Virus or Tracking main effects (*F*<1.1, p>0.2) or interaction of these factors with Choice, (*F*<1.4, *p*>0.3) indicating that for both viral conditions, ST and GT have a similar preference for the non-sated food during choice test.

Figure 4 shows a summary of histological verification and representative examples of viral expression in NAc core (Fig. 4A-B) and BLA (Fig. 4C-D) for hm4di and mCherry constructs. Contralateral injections were counterbalanced, thus for each rat only unilateral cell body expression was observed in contralateral BLA and NAc. Expression is shown in both hemispheres to represent both counterbalanced groups.

**Figure 4.**
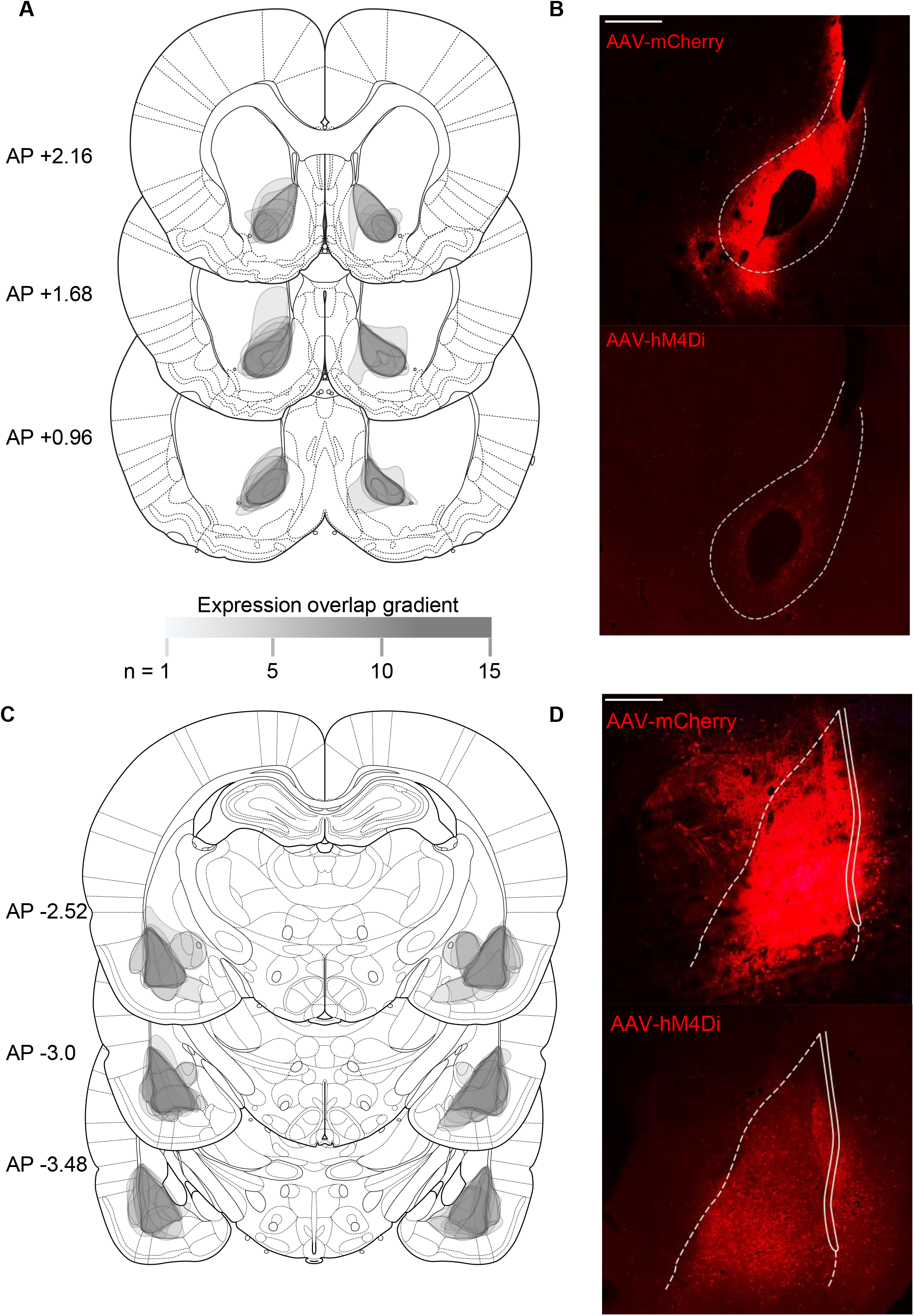
Histological verification of viral expression in NAc core and BLA. Rats were injected with viral constructs unilaterally in BLA and in contralateral NAc core (mm from bregma; (Paxinos & Watson, 2007); scale bars represent 500 μm. Unilateral expression was counterbalanced, but expression is shown in both hemispheres. (A) Schematic representation of viral expression and (B) representative image of mCherry (top) and hM4Di (bottom) NAc core expression. (C) Schematic representation of viral expression and (B) representative image of (top) mCherry and hM4Di (bottom) BLA expression. Legend indicates density of overlapping expression, where (n) is the number of overlapping cases to produce the represented opacity.

## 4 Discussion

We examined the effect of contralaterally inactivating BLA and NAc core on flexibility in outcome devaluation. We found BLA-NAc core inactivation promoted flexibility in otherwise inflexible sign-tracking rats, and disrupted flexibility in otherwise flexible goal-tracking rats. In viral control rats, we replicated previous findings that intact GT rats flexibility reduce approach behavior when the outcome is devalued, while ST rats do not (Keefer et al., 2020; Nasser et al., 2015). The tracking specificity of devaluation sensitivity has been observed across several studies, Pavlovian paradigms, and devaluation procedures (Nasser et al., 2015; Patitucci et al., 2016; Smedley & Smith, 2018; Keefer et al., 2020), but see (Davey & Cleland, 1982; Derman et al., 2018; Amaya et al., 2020). In our study using both males and females, BLA-NAc core contralateral chemogenetic inactivation specifically reduced lever directed behavior, but not food cup-directed behavior, consistent with a prior BLA-NAc crosslesion study showing greater attenuation of lever directed approach in male rats (Chang et al., 2012). While further studies are needed to probe sex differences on the role of BLA-NAc communication in driving devaluation sensitivity, from the present study we predict the tracking-specific effects of this manipulation are carried by male rats.

A body of amygdala lesion and inactivation studies examining the neurobiology of incentive learning (for review see Wassum & Izquierdo, 2015) implicate candidate circuitry that may underlie differences in incentive learning that rely on the motivational properties of cues relative to the current value of the outcome. In brief, pre-training lesions of the BLA impair both the initial acquisition of incentive cue properties as well as subsequent updating of behavior in response to changing outcome values (Hatfield et al., 1996). Post-training lesions of the BLA similarly disrupt behavioral updating during devaluation (Johnson et al., 2009). Additionally, BLA lesions disrupt acquisition of positive incentive value (Setlow, Gallagher, et al., 2002), while lesions of NAc prevent expression of incentive value (McDannald et al., 2013) in SOC, and this pathway is necessary to acquire and express learned motivational value (Setlow, Holland, et al., 2002). Disconnection of the BLA and NAc also produces deficits in both initial acquisition and terminal levels of lever directed behavior, the preferred response of sign-tracking rats (Chang et al., 2012). Thus, we predicted that if ST rats rely on BLA to NAc communication to form rigid, behaviorally inflexible incentive value representations, then inactivation of BLA and NAc core would facilitate behavioral flexibility in outcome devaluation. Consistent with our hypothesis, we observed that ST rats flexibly reduced lever directed behavior during outcome devaluation when BLA and contralateral NAc were inactivated. This suggests that ST rats rely upon these structures to support rigid appetitive approach expressed as lever directed behavior.

Consistent with previous work, we observed that intact GT rats displayed behavioral flexibility, reducing their preferred responding following outcome devaluation, while intact ST rats did not (Morrison et al., 2015; Nasser et al., 2015; Keefer et al., 2020). However, we found GT rats with BLA-NAc chemogenic inactivation were insensitive to devaluation. This finding suggests that GT rats rely upon this circuitry to integrate and/or express learning about changes in reinforcer value. In a PLA task designed to promote goal-tracking responses, NAc core is also necessary for the expression of goal-tracking (Blaiss & Janak, 2009). The present findings are also consistent with prior studies demonstrating that the BLA (Hatfield et al., 1996) and NAc (Singh et al., 2010) are critically involved Pavlovian outcome devaluation. Additionally, disconnection of the BLA and NAc produces a deficit in an instrumental outcome devaluation task (Shiflett & Balleine, 2010). The present study supports the role of this circuit in Pavlovian devaluation and suggests it may support different associative constructs in different individuals. That is, sign-trackers may rely on BLA and NAc to respond to cues based on their appetitive motivational properties, while goal-trackers rely on this circuitry to respond to cues based on the current value of the outcome. Consideration of tracking-specific behavioral and neurobiological differences, as in the present study, may provide a useful framework for interpreting individual variability in circuit manipulation studies.

The tracking-specific role of BLA and NAc core presented here falls into context with prior electrophysiological recording and optogenetic studies. Without BLA excitatory input, NAc fails to represent previously acquired CS-US associations, which blunts conditioned responding directed at both cues and outcomes (Ambroggi et al., 2008; Stuber et al., 2011). Compared to goal-trackers, sign-trackers show attenuated NAc reward signaling and stronger cue-evoked firing as training progresses (Gillis & Morrison, 2019). Similarly, NAc core cue-encoding during second order conditioning positively correlates with SOC performance (Saddoris & Carelli, 2014). Surprisingly, ST and GT rats similarly acquire and express SOC (Saddoris & Carelli, 2014; Nasser et al., 2015), which seems somewhat at odds with the perspective that SOC and ST reflect similar positive incentive learning processes, both of which rely on BLA-NAc communication. Notably, enhanced NAc core cue encoding is also associated with better devaluation performance and sensory preconditioning, two learning processes that reflect an inference about either the current value of the outcome or value-independent predicative stimulus relationships (Cerri et al., 2014; West & Carelli, 2016). The double dissociation we observe in the present study, in which BLA-NAc core inactivation impedes flexibly in ST rats, but facilitates flexibility in GT rats, suggest individual or methodological differences that bias CS or US processing may account for the diverse role for BLA-NAc in inventive learning processes.

### 4.1 Methodological Considerations

Our inclusion of both male and female rats is consistent with current best practices in neuroscience research and is part of a larger, growing trend to improve representation of female subjects in basic science (McCarthy et al., 2017; Miller et al., 2017; Shansky, 2019). For practical reasons we included both males and females without fully powering sex as a factor in order to test our hypothesis about the contribution of BLA and NAc in driving tracking-specific differences in devaluation sensitivity. Consistent with previous work, we observed that females displayed more lever directed behavior than males overall (Madayag et al., 2017 but see Pitchers et al., 2015; Bacharach et al., 2018). Consistent with prior work showing that males are more sensitive to satiety-induced outcome devaluation (Hammerslag & Gulley, 2014), we also see devaluation sensitivity of food cup approach is driven by male rats. While the primary objective of this study was to include both sexes, not to probe sex differences, our exploratory analyses suggest that some sex effects may warrant further investigation. In particular, one testable working hypotheses includes the possibility that the devaluation sensitivity of lever approach that is unmasked by BLA-NAc core inactivation may be sex-specific. The present approach to include and report effects for both sexes ensures we do not rely solely on male rats to determine the causal role of brain circuit contributions to behavior.

The present work does not include the ipsilateral control group that is typical of traditional disconnection designs. In brief, our work employs contralateral chemogenetic inactivation of the BLA and NAc core. To demonstrate that effects are attributable to disrupted BLA-NAc core communication, rather than inactivation of these two regions alone, an ipsilateral control (in which communication between the structures is still possible unilaterally) is often employed. For practical reasons, we were unable to include an ipsilateral control group. However, we are not the first to contralaterally inactivate these regions, and a body of evidence demonstrates no effect of ipsilateral disconnection of the BLA and NAc in similar tasks. Contralateral disconnection of the BLA and NAc disrupts lever-directed approach in Pavlovian lever autoshaping both early and late in training. Critically, ipsilateral controls performed similarly to sham lesioned rats, suggesting unilateral functional communication between BLA and NAc is sufficient to support lever directed behavior (Chang et al., 2012). The present contralateral manipulations replicate the disconnection findings (Chang et al., 2012), bolstering our conjecture that BLA to NAc core communication is what drives our reported effects. Similarly, ipsilateral disconnection of the BLA and NAc produces no impairment in instrumental outcome devaluation or Pavlovian instrumental transfer (Shiflett & Balleine, 2010). Additionally, anatomical evidence establishes BLA to NAc connectivity being primarily unidirectional and unilateral (Swanson & Cowan, 1975; Ottersen, 1980; Russchen & Price, 1984; Heimer et al., 1991; Brog et al., 1993; Kelley et al., 1993). Indeed, excitatory input (either direct or via modulation of dopaminergic inputs) into the NAc originating from the BLA drives neuronal responses to reward-predictive cues (e.g. Floresco et al., 2001; Ambroggi et al., 2008; Simmons & Neil, 2009; Jones et al., 2010). While disconnection of the BLA and NAc reduces neuronal excitability within the NAc and decreases responding toward reward-predictive cues, ipsilateral controls show significantly less pronounced (Ambroggi et al., 2008; muscimol/baclofen inactivation of BLA and D1 antagonism in NAc) or absent changes in excitability and reward-seeking behavior (Simmons & Neil, 2009; muscimol inactivation of BLA and D1/D2 antagonism in NAc). Altogether, while we expect the effects reported here reflect a disruption of communication from BLA to NAc, the ipsilateral control experiments would be necessary to confirm. We conclude that contralateral inactivation of BLA and NAc reveal opposite effects on devaluation sensitivity in sign- and goal-trackers.

### 4.2 Conclusions

Pre-clinical studies evaluating behavioral and neurobiological markers of addiction-vulnerable individuals prior to any drug exposure are an important step toward understanding human addiction. Pre-clinical studies implicate BLA-NAc core communication in driving cocaine seeking (Di Ciano & Everitt, 2004), and NAc is heavily implicated in both sign-tracking and the enhanced cocaine relapse observed in ST rats (Flagel et al., 2011; Chang et al., 2012; Clark et al., 2013; Saunders et al., 2013; Fraser & Janak, 2017;). Sign-trackers show an array of behaviors indicative of maladaptive incentive learning, including resistance to extinction (Ahrens et al., 2016; Fitzpatrick et al., 2019), heightened tolerance for negative consequences (Saunders & Robinson, 2010), and heightened attraction and sensitivity to the reinforcing properties of predictive cues (Flagel et al., 2007; Robinson & Flagel, 2009; Bacharach et al., 2018). While both ST and GT acquire the predictive relationship between cue and reward, ST are thought to attribute a higher level of incentive salience to the cue (Flagel et al., 2009; Pitchers et al., 2015; Flagel & Robinson, 2017). Sign-trackers’ inflexibility prior to and after drug experience (Saunders et al., 2013; Keefer et al., 2020) highlights the utility of the sign-tracking model for understanding the brain basis of SUD vulnerability. This work has translational relevance, as humans also show variability in cue reactivity and devaluation sensitivity (e.g. Garofalo & di Pellegrino, 2015; Versace et al., 2016; De Tommaso et al., 2017; Pool et al., 2019). A deeper understanding of the psychological and neurobiological differences present prior to drug exposure can enhance potential therapeutic interventions (e.g. Saunders & Robinson, 2010, 2013; McClory & Spear, 2014; Versaggi et al., 2016; Pitchers et al., 2017; Valyear et al., 2017). This work also underscores the importance of considering tracking- and sex-specific effects in neurobiological examinations of outcome devaluation. Future studies should be adequately powered to consider sex as a variable, as the present work suggests that there are important sex differences in flexibility that are relevant to addiction vulnerability.

## 6 Data Availability Statement

The raw data supporting the conclusions of this article will be made available by the authors, without undue reservation, to any qualified researcher.

## 7 Ethics Statement

The animal study was reviewed and approved by University of Maryland, School of Medicine Institutional Animal Care and Use Committee.

## 8 Author Contributions

DC conceived and supervised the project. DK, SK, and UG acquired the data. DK analyzed the data. DK and DC designed the experiments, interpreted the data, and wrote the manuscript. All authors contributed to manuscript revision, read, and approved the submitted version.

## 10 Acknowledgements

We thank the UMB Animal Care Facility for colony maintenance. The present affiliation of DK is: Department of Psychology, Washington College, 300 Washington Avenue, Chestertown, MD 21620, USA.

## References

Ahrens, A. M., Singer, B. F., Fitzpatrick, C. J., Morrow, J. D., & Robinson, T. E. (2016). Rats that sign-track are resistant to Pavlovian but not instrumental extinction. Behavioural Brain Research, 296, 418–430. https://doi.org/10.1016/j.bbr.2015.07.055

Amaya, K. A., Stott, J. J., & Smith, K. S. (2020). Sign-tracking behavior is sensitive to outcome devaluation in a devaluation context-dependent manner: Implications for analyzing habitual behavior. Learning & Memory, 27(4), 136–149. https://doi.org/10.1101/lm.051144.119

Ambroggi, F., Ishikawa, A., Fields, H. L., & Nicola, S. M. (2008). Basolateral amygdala neurons facilitate reward-seeking behavior by exciting nucleus accumbens neurons. Neuron, 59(4), 648–661.

Bacharach, S. Z., Nasser, H. M., Zlebnik, N. E., Dantrassy, H. M., Kochli, D. E., Gyawali, U., Cheer, J. F., & Calu, D. J. (2018). Cannabinoid receptor-1 signaling contributions to sign-tracking and conditioned reinforcement in rats. Psychopharmacology, 235(10), 3031–3043. https://doi.org/10.1007/s00213-018-4993-6

Berridge, K. C. (1996). Food reward: Brain substrates of wanting and liking. Neuroscience & Biobehavioral Reviews, 20(1), 1–25. https://doi.org/10.1016/0149-7634(95)00033-B

Berridge, K. C., & Robinson, T. E. (2016). Liking, wanting, and the incentive-sensitization theory of addiction. American Psychologist, 71(8), 670–679. https://doi.org/10.1037/amp0000059

Blaiss, C. A., & Janak, P. H. (2009). The nucleus accumbens core and shell are critical for the expression, but not the consolidation, of Pavlovian conditioned approach. Behavioural brain research, 200(1), 22–32.

Brog, J. S., Salyapongse, A., Deutch, A. Y., & Zahm, D. S. (1993). The patterns of afferent innervation of the core and shell in the “Accumbens” part of the rat ventral striatum: Immunohistochemical detection of retrogradely transported fluoro-gold. Journal of Comparative Neurology, 338(2), 255–278. https://doi.org/10.1002/cne.903380209

Cerri, D. H., Saddoris, M. P., & Carelli, R. M. (2014). Nucleus Accumbens Core Neurons Encode Value-Independent Associations Necessary for Sensory Preconditioning. Behavioral Neuroscience, 128(5), 567–578. https://doi.org/10.1037/a0037797

Chang, S. E., Wheeler, D. S., & Holland, P. C. (2012). Roles of nucleus accumbens and basolateral amygdala in autoshaped lever pressing. Neurobiology of learning and memory, 97(4), 441–451.

Clark, J. J., Collins, A. L., Sanford, C. A., & Phillips, P. E. M. (2013). Dopamine Encoding of Pavlovian Incentive Stimuli Diminishes with Extended Training. The Journal of Neuroscience, 33(8), 3526–3532. https://doi.org/10.1523/JNEUROSCI.5119-12.2013

Cohen, J. (1988). Statistical Power Analysis for the Behavioral Sciences. Lawrence Erlbaum Associates.

Davey, G. C., & Cleland, G. G. (1982). Topography of signal-centered behavior in the rat: Effects of deprivation state and reinforcer type. Journal of the Experimental Analysis of Behavior, 38(3), 291–304. https://doi.org/10.1901/jeab.1982.38-291

Di Ciano, P., & Everitt, B. J. (2004). Direct Interactions between the Basolateral Amygdala and Nucleus Accumbens Core Underlie Cocaine-Seeking Behavior by Rats. Journal of Neuroscience, 24(32), 7167–7173. https://doi.org/10.1523/JNEUROSCI.1581-04.2004

De Tommaso, M., Mastropasqua, T., & Turatto, M. (2017). The salience of a reward cue can outlast reward devaluation. Behavioral Neuroscience, 131(3), 226–234. https://doi.org/10.1037/bne0000193

Derman, R. C., Schneider, K., Juarez, S., & Delamater, A. R. (2018). Sign-tracking is an expectancy-mediated behavior that relies on prediction error mechanisms. Learning & Memory, 25(10), 550–563. https://doi.org/10.1101/lm.047365.118

Fitzpatrick, C. J., Geary, T., Creeden, J. F., & Morrow, J. D. (2019). Sign-tracking behavior is difficult to extinguish and resistant to multiple cognitive enhancers. Neurobiology of Learning and Memory, 163, 107045. https://doi.org/10.1016/j.nlm.2019.107045

Flagel, S. B., Akil, H., & Robinson, T. E. (2009). Individual differences in the attribution of incentive salience to reward-related cues: Implications for addiction. Neuropharmacology, 56, 139–148. https://doi.org/10.1016/j.neuropharm.2008.06.027

Flagel, S. B., Clark, J. J., Robinson, T. E., Mayo, L., Czuj, A., Willuhn, I., Akers, C. A., Clinton, S. M., Phillips, P. E. M., & Akil, H. (2011). A selective role for dopamine in reward learning. Nature, 469(7328), 53–57. https://doi.org/10.1038/nature09588

Flagel, S. B., & Robinson, T. E. (2017). Neurobiological basis of individual variation in stimulus-reward learning. Current Opinion in Behavioral Sciences, 13, 178–185. https://doi.org/10.1016/j.cobeha.2016.12.004

Flagel, S. B., Watson, S. J., Robinson, T. E., & Akil, H. (2007). Individual differences in the propensity to approach signals vs goals promote different adaptations in the dopamine system of rats. Psychopharmacology, 191(3), 599–607. https://doi.org/10.1007/s00213-006-0535-8

Floresco, S. B., Blaha, C. D., Yang, C. R., & Phillips, A. G. (2001). Dopamine D1 and NMDA receptors mediate potentiation of basolateral amygdala-evoked firing of nucleus accumbens neurons. Journal of Neuroscience, 21(16), 6370–6376.

Fraser, K. M., & Janak, P. H. (2017). Long-lasting contribution of dopamine in the nucleus accumbens core, but not dorsal lateral striatum, to sign-tracking. The European Journal of Neuroscience, 46(4), 2047–2055. https://doi.org/10.1111/ejn.13642

Garofalo, S., & di Pellegrino, G. (2015). Individual differences in the influence of task-irrelevant Pavlovian cues on human behavior. Frontiers in Behavioral Neuroscience, 9. https://doi.org/10.3389/fnbeh.2015.00163

Gillis, Z. S., & Morrison, S. E. (2019). Sign Tracking and Goal Tracking Are Characterized by Distinct Patterns of Nucleus Accumbens Activity. ENeuro, 6(2). https://doi.org/10.1523/ENEURO.0414-18.2019

Gomez, J. L., Bonaventura, J., Lesniak, W., Mathews, W. B., Sysa-Shah, P., Rodriguez, L. A., … & Pomper, M. G. (2017). Chemogenetics revealed: DREADD occupancy and activation via converted clozapine. Science, 357(6350), 503–507.

Hammerslag, L. R., & Gulley, J. M. (2014). Age and sex differences in reward behavior in adolescent and adult rats. Developmental Psychobiology, 56(4), 611–621. https://doi.org/10.1002/dev.21127

Hatfield, T., Han, J.-S., Conley, M., Gallagher, M., & Holland, P. (1996). Neurotoxic Lesions of Basolateral, But Not Central, Amygdala Interfere with Pavlovian Second-Order Conditioning and Reinforcer Devaluation Effects. Journal of Neuroscience, 16(16), 5256–5265. https://doi.org/10.1523/JNEUROSCI.16-16-05256.1996

Heimer, L., Zahm, D. S., Churchill, L., Kalivas, P. W., & Wohltmann, C. (1991). Specificity in the projection patterns of accumbal core and shell in the rat. Neuroscience, 41(1), 89–125. https://doi.org/10.1016/0306-4522(91)90202-Y

Holland, P. C., & Rescorla, R. A. (1975). Second-order conditioning with food unconditioned stimulus. Journal of Comparative and Physiological Psychology, 88(1), 459–467. https://doi.org/10.1037/h0076219

Johnson, A. W., Gallagher, M., & Holland, P. C. (2009). The Basolateral Amygdala Is Critical to the Expression of Pavlovian and Instrumental Outcome-Specific Reinforcer Devaluation Effects. The Journal of Neuroscience, 29(3), 696–704. https://doi.org/10.1523/JNEUROSCI.3758-08.2009

Kawa, A. B., Bentzley, B. S., & Robinson, T. E. (2016). Less is more: Prolonged intermittent access cocaine self-administration produces incentive-sensitization and addiction-like behavior. Psychopharmacology, 233(19–20), 3587–3602. https://doi.org/10.1007/s00213-016-4393-8

Keefer, S. E., Bacharach, S. Z., Kochli, D. E., Chabot, J. M., & Calu, D. J. (2020). Effects of Limited and Extended Pavlovian Training on Devaluation Sensitivity of Sign- and Goal-Tracking Rats. Frontiers in Behavioral Neuroscience, 14. https://doi.org/10.3389/fnbeh.2020.00003

Kelley, A. E., Domesick, V. B., & Nauta, W. J. H. (1993). The Amygdalostriatal Projection in the Rat—An Anatomical Study by Anterograde and Retrograde Tracing Methods. In W. J. H. Nauta (Ed.), Neuroanatomy (pp. 495–509). Birkhäuser. https://doi.org/10.1007/978-1-4684-7920-1_24

Madayag, A. C., Stringfield, S. J., Reissner, K. J., Boettiger, C. A., & Robinson, D. L. (2017). Sex and Adolescent Ethanol Exposure Influence Pavlovian Conditioned Approach. Alcoholism: Clinical and Experimental Research, 41(4), 846–856. https://doi.org/10.1111/acer.13354

McCarthy, M. M., Woolley, C. S., & Arnold, A. P. (2017). Incorporating sex as a biological variable in neuroscience: What do we gain? Nature Reviews Neuroscience, 18(12), 707–708. https://doi.org/10.1038/nrn.2017.137

McClory, A. J., & Spear, L. P. (2014). Effects of ethanol exposure during adolescence or in adulthood on Pavlovian conditioned approach in Sprague-Dawley rats. Alcohol, 48(8), 755–763. https://doi.org/10.1016/j.alcohol.2014.05.006

McDannald, M. A., Setlow, B., & Holland, P. C. (2013). Effects of ventral striatal lesions on first- and second-order appetitive conditioning. European Journal of Neuroscience, 38(4), 2589–2599. https://doi.org/10.1111/ejn.12255

Meyer, P. J., Lovic, V., Saunders, B. T., Yager, L. M., Flagel, S. B., Morrow, J. D., & Robinson, T. E. (2012). Quantifying Individual Variation in the Propensity to Attribute Incentive Salience to Reward Cues. PLoS ONE, 7(6). https://doi.org/10.1371/journal.pone.0038987

Miller, L. R., Marks, C., Becker, J. B., Hurn, P. D., Chen, W.-J., Woodruff, T., McCarthy, M. M., Sohrabji, F., Schiebinger, L., Wetherington, C. L., Makris, S., Arnold, A. P., Einstein, G., Miller, V. M., Sandberg, K., Maier, S., Cornelison, T. L., & Clayton, J. A. (2017). Considering sex as a biological variable in preclinical research. The FASEB Journal, 31(1), 29–34. https://doi.org/10.1096/fj.201600781r

Morrison, S. E., Bamkole, M. A., & Nicola, S. M. (2015). Sign Tracking, but Not Goal Tracking, is Resistant to Outcome Devaluation. Frontiers in Neuroscience, 9. https://doi.org/10.3389/fnins.2015.00468

Nasser, H. M., Chen, Y.-W., Fiscella, K., & Calu, D. J. (2015). Individual variability in behavioral flexibility predicts sign-tracking tendency. Frontiers in Behavioral Neuroscience, 9. https://doi.org/10.3389/fnbeh.2015.00289

Ottersen, O. P. (1980). Afferent connections to the amygdaloid complex of the rat and cat: II. Afferents from the hypothalamus and the basal telencephalon. Journal of Comparative Neurology, 194(1), 267–289. https://doi.org/10.1002/cne.901940113

Parkes, S. L., & Balleine, B. W. (2013). Incentive memory: evidence the basolateral amygdala encodes and the insular cortex retrieves outcome values to guide choice between goal-directed actions. Journal of Neuroscience, 33(20), 8753–8763.

Patitucci, E., Nelson, A. J. D., Dwyer, D. M., & Honey, R. C. (2016). The origins of individual differences in how learning is expressed in rats: A general-process perspective. Journal of Experimental Psychology: Animal Learning and Cognition, 42(4), 313. https://doi.org/10.1037/xan0000116

Paxinos, G., & Watson, C. (2007). The rat brain in stereotaxic coordinates in stereotaxic coordinates. Elsevier.

Pitchers, K. K., Flagel, S. B., O’Donnell, E. G., Solberg Woods, L. C., Sarter, M., & Robinson, T. E. (2015). Individual variation in the propensity to attribute incentive salience to a food cue: Influence of sex. Behavioural Brain Research, 278, 462–469. https://doi.org/10.1016/j.bbr.2014.10.036

Pitchers, K. K., Phillips, K. B., Jones, J. L., Robinson, T. E., & Sarter, M. (2017). Diverse Roads to Relapse: A Discriminative Cue Signaling Cocaine Availability Is More Effective in Renewing Cocaine Seeking in Goal Trackers Than Sign Trackers and Depends on Basal Forebrain Cholinergic Activity. Journal of Neuroscience, 37(30), 7198–7208. https://doi.org/10.1523/JNEUROSCI.0990-17.2017

Pool, E. R., Pauli, W. M., Kress, C. S., & O’Doherty, J. P. (2019). Behavioural evidence for parallel outcome-sensitive and outcome-insensitive Pavlovian learning systems in humans. Nature Human Behaviour, 3(3), 284–296. https://doi.org/10.1038/s41562-018-0527-9

Robinson, T. E., & Berridge, K. C. (1993). The neural basis of drug craving: An incentive-sensitization theory of addiction. Brain Research Reviews, 18(3), 247–291. https://doi.org/10.1016/0165-0173(93)90013-P

Robinson, T. E., & Flagel, S. B. (2009). Dissociating the Predictive and Incentive Motivational Properties of Reward-Related Cues Through the Study of Individual Differences. Biological Psychiatry, 65(10), 869–873. https://doi.org/10.1016/j.biopsych.2008.09.006

Rode, A. N., Moghaddam, B., & Morrison, S. E. (2020). Increased Goal Tracking in Adolescent Rats Is Goal-Directed and Not Habit-Like. Frontiers in Behavioral Neuroscience, 13. https://doi.org/10.3389/fnbeh.2019.00291

Russchen, F. T., & Price, J. L. (1984). Amygdalostriatal projections in the rat. Topographical organization and fiber morphology shown using the lectin PHA-L as an anterograde tracer. Neuroscience Letters, 47(1), 15–22. https://doi.org/10.1016/0304-3940(84)90379-3

Saddoris, M. P., & Carelli, R. M. (2014). Cocaine Self-Administration Abolishes Associative Neural Encoding in the Nucleus Accumbens Necessary for Higher-Order Learning. Biological Psychiatry, 75(2). https://doi.org/10.1016/j.biopsych.2013.07.037

Saddoris, M. P., Wang, X., Sugam, J. A., & Carelli, R. M. (2016). Cocaine Self-Administration Experience Induces Pathological Phasic Accumbens Dopamine Signals and Abnormal Incentive Behaviors in Drug-Abstinent Rats. The Journal of Neuroscience, 36(1), 235–250. https://doi.org/10.1523/JNEUROSCI.3468-15.2016

Saunders, B. T., & Robinson, T. E. (2010). A Cocaine Cue Acts as an Incentive Stimulus in Some but not Others: Implications for Addiction. Biological Psychiatry, 67(8), 730–736. https://doi.org/10.1016/j.biopsych.2009.11.015

Saunders, B. T., & Robinson, T. E. (2013). Individual variation in resisting temptation: Implications for addiction. Neuroscience & Biobehavioral Reviews, 37(9, Part A), 1955–1975. https://doi.org/10.1016/j.neubiorev.2013.02.008

Saunders, B. T., Yager, L. M., & Robinson, T. E. (2013). Cue-Evoked Cocaine “Craving”: Role of Dopamine in the Accumbens Core. Journal of Neuroscience, 33(35), 13989–14000. https://doi.org/10.1523/JNEUROSCI.0450-13.2013

Setlow, B., Gallagher, M., & Holland, P. C. (2002). The basolateral complex of the amygdala is necessary for acquisition but not expression of CS motivational value in appetitive Pavlovian second-order conditioning. European Journal of Neuroscience, 15(11), 1841–1853. https://doi.org/10.1046/j.1460-9568.2002.02010.x

Setlow, B., Holland, P. C., & Gallagher, M. (2002). Disconnection of the basolateral amygdala complex and nucleus accumbens impairs appetitive Pavlovian second-order conditioned responses. Behavioral Neuroscience, 116(2), 267–275. https://doi.org/10.1037/0735-7044.116.2.267

Shansky, R. M. (2019). Are hormones a “female problem” for animal research?. Science, 364(6443), 825–826.

Shiflett, M. W., & Balleine, B. W. (2010). At the limbic–motor interface: Disconnection of basolateral amygdala from nucleus accumbens core and shell reveals dissociable components of incentive motivation. European Journal of Neuroscience, 32(10), 1735–1743. https://doi.org/10.1111/j.1460-9568.2010.07439.x

Simmons, D. A., & Neill, D. B. (2009). Functional interaction between the basolateral amygdala and the nucleus accumbens underlies incentive motivation for food reward on a fixed ratio schedule. Neuroscience, 159(4), 1264–1273.

Singh, T., McDannald, M., Haney, R., Cerri, D., & Schoenbaum, G. (2010). Nucleus Accumbens Core and Shell are Necessary for Reinforcer Devaluation Effects on Pavlovian Conditioned Responding. Frontiers in Integrative Neuroscience, 4. https://doi.org/10.3389/fnint.2010.00126

Smedley, E. B., & Smith, K. S. (2018). Evidence of structure and persistence in motivational attraction to serial Pavlovian cues. Learning & Memory, 25(2), 78–89. https://doi.org/10.1101/lm.046599.117

Stuber, G. D., Sparta, D. R., Stamatakis, A. M., van Leeuwen, W. A., Hardjoprajitno, J. E., Cho, S., Tye, K. M., Kempadoo, K. A., Zhang, F., Deisseroth, K., & Bonci, A. (2011). Amygdala to nucleus accumbens excitatory transmission facilitates reward seeking. Nature, 475(7356), 377–380. https://doi.org/10.1038/nature10194

Swanson, L. W., & Cowan, W. M. (1975). A note on the connections and development of the nucleus accumbens. Brain Research, 92(2), 324–330. https://doi.org/10.1016/0006-8993(75)90278-4

Tomie, A., Grimes, K. L., & Pohorecky, L. A. (2008). Behavioral characteristics and neurobiological substrates shared by Pavlovian sign-tracking and drug abuse. Brain Research Reviews, 58(1), 121–135. https://doi.org/10.1016/j.brainresrev.2007.12.003

Valyear, M. D., Villaruel, F. R., & Chaudhri, N. (2017). Alcohol-seeking and relapse: A focus on incentive salience and contextual conditioning. Behavioural Processes, 141, 26–32. https://doi.org/10.1016/j.beproc.2017.04.019

Versace, F., Kypriotakis, G., Basen-Engquist, K., & Schembre, S. M. (2016). Heterogeneity in brain reactivity to pleasant and food cues: Evidence of sign-tracking in humans. Social Cognitive and Affective Neuroscience, 11(4), 604–611. https://doi.org/10.1093/scan/nsv143

Versaggi, C. L., King, C. P., & Meyer, P. J. (2016). The tendency to sign-track predicts cue-induced reinstatement during nicotine self-administration, and is enhanced by nicotine but not ethanol. Psychopharmacology, 233(15), 2985–2997. https://doi.org/10.1007/s00213-016-4341-7

Villaruel, F. R., & Chaudhri, N. (2016). Individual Differences in the Attribution of Incentive Salience to a Pavlovian Alcohol Cue. Frontiers in Behavioral Neuroscience, 10. https://doi.org/10.3389/fnbeh.2016.00238

Wassum, K. M., & Izquierdo, A. (2015). The basolateral amygdala in reward learning and addiction. Neuroscience & Biobehavioral Reviews, 57, 271–283. https://doi.org/10.1016/j.neubiorev.2015.08.017

West, E. A., & Carelli, R. M. (2016). Nucleus Accumbens Core and Shell Differentially Encode Reward-Associated Cues after Reinforcer Devaluation. The Journal of Neuroscience, 36(4), 1128–1139. https://doi.org/10.1523/JNEUROSCI.2976-15.2016

Yager, L. M., Pitchers, K. K., Flagel, S. B., & Robinson, T. E. (2015). Individual Variation in the Motivational and Neurobiological Effects of an Opioid Cue. Neuropsychopharmacology, 40(5), 1269–1277. https://doi.org/10.1038/npp.2014.314

